# Viral diversity is an obligate consideration in CRISPR/Cas9 designs for HIV cure

**DOI:** 10.1101/255133

**Authors:** Pavitra Roychoudhury, Harshana De Silva Feelixge, Daniel Reeves, Bryan T. Mayer, Daniel Stone, Joshua T. Schiffer, Keith R. Jerome

## Abstract

RNA-guided CRISPR/Cas9 systems can be designed to mutate or excise the integrated HIV genome from latently infected cells and have therefore been proposed as a curative approach for HIV. However, most studies to date have focused on molecular clones with ideal target site recognition and do not account for target site variability observed within and between patients. For clinical success and broad applicability, guide RNA (gRNA) selection must account for circulating strain diversity and incorporate the within-host diversity of HIV. To address this, we identified a set of gRNAs targeting HIV LTR, *gag* and *pol* using publicly available sequences for these genes. We ranked gRNAs according to global conservation across HIV-1 group M and within subtypes A-C. By considering paired and triplet combinations of gRNAs, we found triplet sets of target sites such that at least one of the gRNAs in the set was present in over 98% of all globally-available sequences. We then selected 59 gRNAs from our list of highly-conserved LTR target sites and evaluated *in vitro* activity using a loss-of-function LTR-GFP fusion reporter. We achieved efficient GFP knockdown with multiple gRNAs and found clustering of highly active gRNA target sites near the middle of the LTR. Using published deep-sequence data from HIV-infected patients, we found that globally conserved sites also had greater within-host target conservation. Lastly, we developed a mathematical model based on varying distributions of within-host HIV sequence diversity and enzyme efficacy. We used the model to estimate the number of doses required to deplete the latent reservoir and achieve functional cure thresholds. Our modeling results highlight the importance of within-host target site conservation. While increased doses may overcome low target cleavage efficiency, inadequate targeting of rare strains is predicted to lead to rebound upon ART cessation even with many doses.

**Author summary:** The field of genome engineering has exploded over the last decade with the discovery of targeted endonucleases such as CRISPR/Cas9. Endonucleases are now being used to develop a wide range of therapeutics and their use has expanded into antiviral therapy against latent viral infections like HIV. The idea is to induce mutations in latent viral genomes that will render them replication-incompetent, thereby producing a functional cure. Although a great deal of progress has been made, most studies to date have relied on molecular clones that represent “ideal” targets. For clinical success and broad applicability, these therapies need to account for viral genetic diversity within and between individuals. Our paper examines the impact of HIV diversity on CRISPR-based cure strategies to determine the predictors of future clinical success. We performed an exhaustive and detailed computational analysis to identify optimal CRISPR target sites, taking into consideration both within-host and global viral diversity. We coupled this with laboratory testing of highly-conserved guides and compared measured activity to predicted results. Finally, we developed a mathematical model to predict the impact of enzyme activity and viral diversity on the number of doses of a CRISPR-based therapy needed to achieve a functional cure of HIV.

## Introduction

Despite the success of combination antiretroviral therapy (cART) in suppressing HIV viremia, reservoirs of latently-infected cells remain the major barrier for HIV cure [1]. The HIV latent reservoir is composed of long-lived infected cells harboring replication-competent proviruses with limited transcription that can reactivate and reseed the reservoir upon cART interruption [2,3]. A promising therapeutic strategy for achieving cure involves depleting the reservoir by direct disruption of proviral genomes using engineered DNA-editing enzymes such as CRISPR/Cas9 nucleases. A growing body of research shows that endonuclease-induced mutation of essential viral genes or excision of provirus can render the virus unable to replicate [4–12]. If performed on a large scale, this approach could yield pharmacologically significant reservoir reduction. However, viral reservoirs are highly diverse, even in well-suppressed individuals [13,14], and this diversity remains a major challenge for the application of genome editing strategies towards an HIV cure. Effective targeting of all viral genetic variants within an infected individual will be crucial for achieving sufficient reservoir reduction to prevent viral rebound upon cART cessation [15,16] and preventing the emergence of resistance to this therapy [11].

Thus far, studies used to demonstrate the viability of gene editing strategies against HIV have primarily targeted single molecular clones that provide ideal endonuclease target site recognition [7,8]. Multiple classes of gene-editing enzymes have been studied, but the CRISPR/Cas9 system has gained popularity in recent years due to its effectiveness, relative simplicity, and ease of use. Several computational tools now exist to identify CRISPR target sites, to predict the activity of guide RNAs (gRNAs) targeting those sites, and to identify and score gRNAs based on multiple factors including predicted off-target activity [17–19]. However, no available tools allow guide selection based on predicted target site conservation or predicted clinical efficacy based on viral diversity. The identification and characterization of the most conserved target sites on a group- or subtype-specific basis will allow rapid selection of gRNAs when deep sequencing of a patient’s reservoir is not practical or feasible. Furthermore, because the virus can evolve resistance to endonuclease targeting [11], multiple sites may need to be targeted concurrently in order to prevent the emergence of resistance. Therefore, the selection of multiplexed sets of gRNAs must account for the diversity of circulating strains across a wide range of infected people, and dosing strategies must consider within-host diversity of HIV to maximize the probability of a functional cure.

Here we present a CRISPR gRNA design strategy that selects target sites not only by predicted efficacy and specificity but also by prevalence in the population. We first created a database of highly conserved target sites in HIV LTR, *gag*, and *pol* focusing on group- and subtype-level conservation using information about the global sequence diversity of HIV. We used this database to identify highly-conserved target site pairs and triplets to create multiplex gRNA designs predicted to maximize targeting and reduce the probability of treatment resistance. From this analysis, we identified and tested 59 LTR guides using a fluorescent reporter to quantify activity *in vitro*. We then used deep-sequence data from HIV-infected individuals to determine target site conservation and probability of cleavage by individual gRNAs in our list. Finally, we used a mathematical model to predict the number of doses that would be required to achieve functional cure thresholds, while accounting for varying levels of target site diversity and enzyme efficacy.

## Results

### Broadly-targeting spCas9 gRNAs against HIV gag, pol, and LTR

We performed a screen to identify globally conserved target sites for *Streptococcus pyogenes* (spCas9) in LTR, *gag*, and *pol* using alignments for these regions obtained from the HIV LANL database. LTR was chosen for its utility in excision of the provirus [20–22], while *gag* and *pol* were chosen based on their conservation between HIV strains [23]. The publicly-available LANL alignments contain HIV sequences from thousands of infected persons (from about 1200 for LTR to more than 8000 for *pol*) and include strain and geographic information. From these alignments, we computed majority consensus sequences for LTR, *gag*, and *pol* of HIV-1 group M and subtypes A-C. We identified a total of 246 unique gRNA target sites in LTR, 573 in *gag*, and 897 in *pol*. For each target site identified, we determined the number of exact hits in the overall alignment of all group M sequences and for each subtype, and ranked target sites by overall prevalence (Fig 1). Target sites were found to be most conserved in *pol* (Table 1), where a single target site was present in up to 86.5% (n = 4416) of all group M sequences. The most-conserved target sites in LTR and *gag* occurred in up to 70.6% (n = 1216) and 71.1% (n = 8435) of group M sequences respectively.

**Fig 1.**
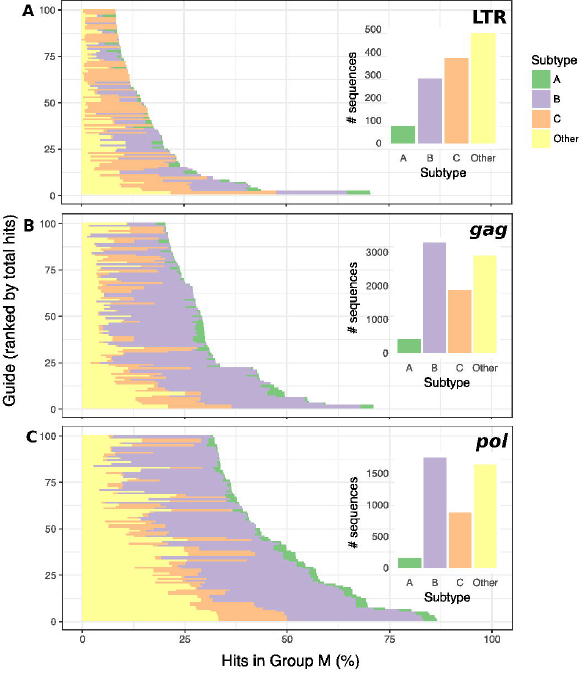
Top 100 gRNA target sites in HIV LTR (A), *gag* (B) and *pol* (C) ranked by prevalence (bottom to top) within an alignment of available sequences within group M for each genomic region. The x-axis shows the percentage of all sequences in group M that contain an exact match to the target site. Within each horizontal bar, shading indicates what percentage of sequences with target sites hits belong to each subtype. Inset bar plots show the total number of sequences of each subtype in the alignment.

**Table 1.**
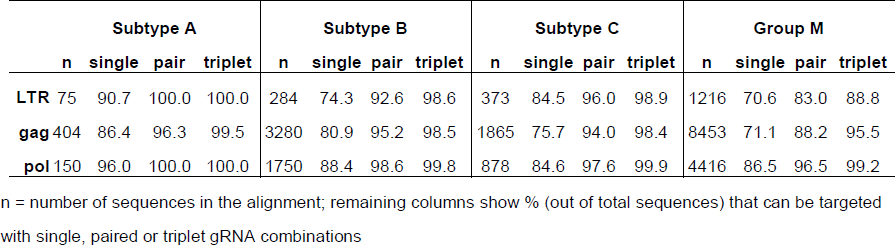
**Maximum targeting possible with 1, 2 or 3 gRNAs**

We determined predicted on-target cleavage efficiency and off-target activity for each guide sequence (Fig 2) using the sgRNA designer tool [17]. Predicted on-target activity scores were in the range [0,1] where a score of 1 was associated with successful knockout in the experiments of Doench et al. [17,24] and gRNAs with scores < 0.2 were generally excluded because they were shown to be predictive of poor activity. Mean predicted activity scores across all identified guides were 0.50 (SD 0.12, n = 246) for LTR, 0.49 (SD 0.13, n = 573) for *gag* and 0.47 (SD 0.13, n = 897) for *pol*. From the list of gRNAs identified, we excluded 10 from *gag* and 26 from *pol* from further analyses due to high predicted off-target activity scores. No significant correlation was observed between predicted activity and target site conservation (Table S1A).

**Fig 2.**
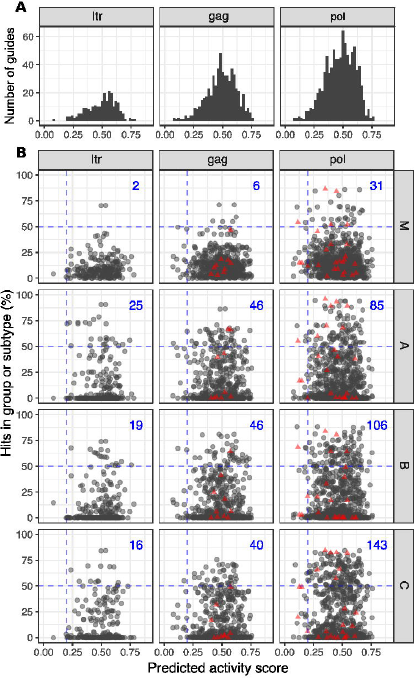
(A) Histogram of predicted activity of all gRNAs identified in LTR, *gag* and *pol* across all four consensus sequences (group M, subtype A-C) for each gene. (B) Predicted activity score vs. target site conservation for individual gRNAs grouped by subtype and gene. Red triangles indicate gRNAs excluded due to predicted off-target activity. Numbers in blue represent the total number of guides with predicted activity score > 0.2 and where target sites occur in more than 50% of sequences in the group or subtype alignment.

### Multiplexed gRNA designs

For each gene, we determined the number of sequences that could be targeted by pairs and triplets of gRNAs in group M overall, and in each subtype A-C. We determined that just 2 strategically selected gRNAs are sufficient for targeting 100% of LTR and *pol* sequences in the current global alignment for Subtype A, and 3 gRNAs are able to target over 98% of all sequences in Subtypes A-C. However, when considering all group M sequences, the maximum percentage of sequences targeted by triplet sets of gRNAs drops to 88.8% for LTR, 95.5% for *gag* and 99.2% for *pol* (Table 1). Overall, better coverage of group M, or subtypes A-C sequences was achieved when pair or triplet gRNAs targeted *pol* suggesting that *pol* is an ideal therapeutic target for targeted mutagenesis with multiplexed guide RNAs. The two most conserved LTR sites in the whole of group M (rank 1 and 2) were also the most prevalent target sites in the individual subtypes, but this was not the case for *gag* and *pol* (Table S2).

### Functional testing of selected gRNAs

From our list of 246 gRNAs targeting LTR, we identified 59 gRNAs for functional testing by first considering the most conserved target sites in group M and each subtype. We then included any gRNAs that increase the number of sequences targeted when combined in pairs or triplets with the previous list (Fig S1A). In order to test the activity of these guides in vitro, we designed LTR-GFP fusion reporter constructs using consensus sequences for group M and subtypes A-C (Fig 3A, Fig S1B). We tested the ability of each gRNA to knock down reporter GFP expression in HEK293 cells following co-transfection with a plasmid expressing spCas9 mCherry containing each HIV-specific gRNA and the LTR-GFP fusion reporter. The activity of each gRNA was measured in terms of percent knockdown of median GFP fluorescence intensity relative to negative controls at 24 h post-transfection in Cas9 expressing (mCherry positive, Fig S1C) cells.

**Fig 3.**
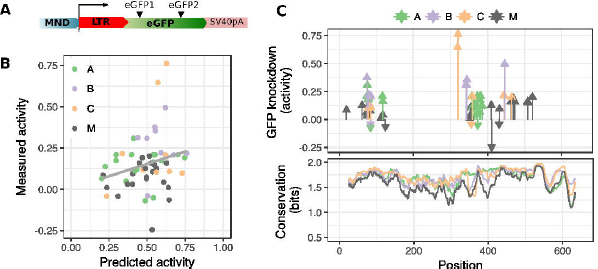
(A) LTR-GFP fusion reporter to test gRNAs for activity *in vitro*. (B) Activity was measured in terms of % knockdown of median GFP fluorescence intensity relative to negative controls. We found positive but statistically non-significant correlation between computationally predicted activity scores and measured activity. (C) We achieved reduction of GFP fluorescence intensity (positive activity) with a majority of gRNA designs and observed clustering of tested target sites in two areas of the LTR with the most active guides being clustered around the center of the LTR. With a small number of gRNAs, we observed negative activity (increase in GFP fluorescence). Lower panel shows residue conservation (in 0-2 bits) across the LTR for alignments of subtype sequences or all sequences in group M.

We compared measured gRNA activity to predicted activity scores from the sgRNA designer (Fig 3B); there was a trend towards weak positive correlation between predicted and measured activity (Pearson’s r = 0.25, n = 59, 95% CI = 0.00–0.48, Table S1B). We observed a reduction of GFP fluorescence intensity with 52 out of 59 gRNAs (Fig 3C, Table S3), with a maximum knockdown of 76.3% (mean = 15.3%, SD = 16.0%, n = 59). Maximum knockdown was achieved at target site CAAAGACTGCTGACACAGAAGGG, which was identified in the consensus sequence of subtype C and found to occur in 23.1% of group M sequences and 68.4% of subtype C sequences in the 2016 LANL alignment. We observed clustering of the most active guides within the LTR; target sites for gRNAs with GFP knockdown > 30% were found at positions 74-75, 319-344 and 446 relative to the start of the 5’ LTR. Although some active guides appear to coincide with regions of high residue conservation within the LTR (Fig 3C), we found no significant correlation between GFP knockdown and target site prevalence within all available sequences in Group M (Pearson’s r = -0.03, n = 59, 95% CI = -0.28–0.23, Table S1C).

### In silico testing of candidate gRNAs on within-host patient sequences

In order to simulate the application of this gene-editing approach on a diverse within-host virus population, we used a published dataset of HIV sequences obtained from HIV-infected blood donors in Brazil [25], focusing on the *pol* gene (because it is the most highly conserved) for 10 patients. We started with our list of all *pol* target sites that we identified above from group and subtype consensus sequences from 2016 LANL alignments, labelling each target site according to the consensus sequence it was identified from (300, 317, 304 and 328 target sites from group M and subtype A-C consensus sequences respectively, 1249 sites total, 897 unique sites). From this combined list of globally conserved target sites, we determined whether each site was present in each patient’s HIV consensus sequence (Tables S4 and S5) [25]. Across infected persons, an average of 89.4 group M target sites (i.e. 29.80% of all group M sites identified) and 119.9 subtype B sites (39.44% of all subtype B target sites identified) were found to be also present within patient consensus sequences (SD 11.14 sites/3.24% and 9.84 sites/3.71% respectively, n = 10 patients), while subtype A and C sites were identified less frequently (Fig 4A). Since subtype B is highly prevalent in Brazil this was not surprising. Five target sites were found to be present in all 10 patient consensus sequences (Table S5) and one of these (GATGGCAGGTGATGATTGTGTGG) was also highly conserved in the global alignment for subtype B (present in 87.09% of LANL sequences). These five target sites were found to occur between positions 2294 and 2981 in *pol*. In addition, we identified gRNA target sites directly from the patient’s consensus sequence. The number of directly identified sites for each patient ranged between 276 and 313 (mean = 299.30, SD = 10.83, n = 10). Out of 1712 unique sites generated from the 10 patients’ consensus sequences, 351 were present in our list of globally conserved sites. Of the remaining sites, 1135 were only present in a single individual and 87 223 sites were found in more than 5 individuals. With one exception (GTTTCTTGCCCTGTCTCTGCTGG), every site that was present in more than 5 individuals was also present in our global list.

**Fig 4.**
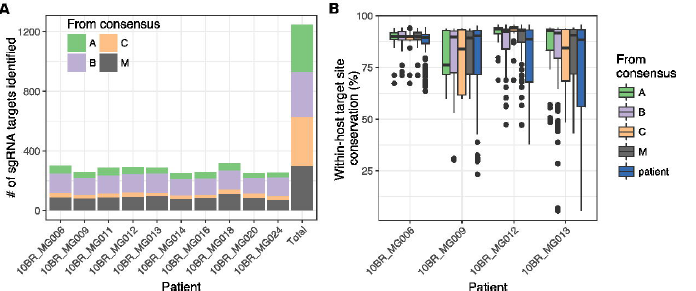
(A) Number of previously identified target sites from global consensus sequences of group M and subtype A-C that were present in each patient’s HIV consensus sequence. (B) Within-host target site conservation for each identified target site using deep-sequence data for 4 patients, summarized using box plots. Black dots indicate outlier target sites (outside 1.5xIQR) and target sites are grouped and colored according to which consensus sequence they were identified from (the group- or subtype-level consensus from LANL alignments, or from the patient’s HIV consensus sequence).

Next we used deep-sequence data from each of these individuals [25] to determine the degree of conservation of each target site within the patient’s virus quasispecies population. In order to accurately quantify rare target site variants, we identified 4 out of 10 patient datasets where mean coverage across all identified target sites was above 5000x (Table S2, Fig S2B). For each of these patients, we determined within-host target site conservation by computing the percentage of reads in the alignment containing an exact match to the site. Within-host target site conservation was found to vary dramatically for individual gRNAs and between individual patients, ranging between 5.5% and 95.6% with a mean of 83.5% (SD 14.3%, n = 2298) (Fig 4B).

Within-host target site conservation was an average of 3.4% higher for sites identified from our global list (range of means = 84.7% - 86.5%, n = 4 patients) compared to sites that were only present in the patient’s sequence (mean = 81.6%, n = 4, p = 0.026) but the difference between groups was not statistically significant (F-test, p = 0.15). Target sites identified from group M or subtype B consensus sequences tended to be more conserved than sites identified from the patient sequence, but the differences were not statistically significant (both 3.7% higher, with p = 0.087 and p = 0.054, respectively). Within-host target site conservation was nearly identical using group M or subtype B sites (p = 0.98). All p-values were > 0.1 after multiple test corrections.

### Modelling reservoir depletion with CRISPR-based therapy

We developed a mathematical model to understand the effect of experimentally-controllable parameters on reservoir depletion with hypothetical weekly dosing of various candidate CRISPR/Cas9 therapies targeting HIV. The model simulates the decay of the latent reservoir by including many (up to 10^4^) quasispecies carrying replication-competent DNA. These species are unevenly abundant, and are assumed to follow a log-normal distribution so that each quasispecies contains 1-1,000 members. Further, each quasispecies decays in size with a rate drawn from the distribution of reservoir decay rates (half-life 3-4 years) described previously [26,27]. In the absence of CRISPR therapy, the model simulates a fluctuating but, on average, slowly decaying HIV reservoir with varying compositions [28]. The parameters analyzed were enzyme efficacy (ϵ, the probability of successful mutagenic DNA cleavage at the target site) and coverage proportion (ρ, the proportion of sequences that would respond to enzyme). The measure of target site conservation is based on our analysis of patient samples.

Including CRISPR, our simulations suggest that treatments with gRNAs targeting a single site will be insufficient to achieve functional cure even at high levels of target site conservation (99%) and enzyme efficiency (0.99) (Fig 5A, Fig S3). Enzyme efficacy is relatively unimportant in this case, only affecting the number of treatments needed to remove the sensitive quasispecies. Once removed, additional treatments provide no additional benefit because insensitive quasispecies dominate the reservoir (Fig 5B). However, if it is possible to achieve 100% coverage of all quasispecies through the selection of a multiplexed set of gRNAs that can be delivered simultaneously, the number of treatments to deplete the reservoir to the first cure threshold (100-fold decrease [16]) can be achieved in 1-5 treatments depending on efficacy (Fig 5C), whereas the second threshold (10^4^-fold decrease [15]) requires 10-15 treatments depending on efficacy. For all modeled assumptions, coverage is vital to reservoir depletion. Whereas suboptimal efficiency can be surmounted by repeated doses, the diversity of the reservoir provides the largest barrier to depletion.

**Fig 5.**
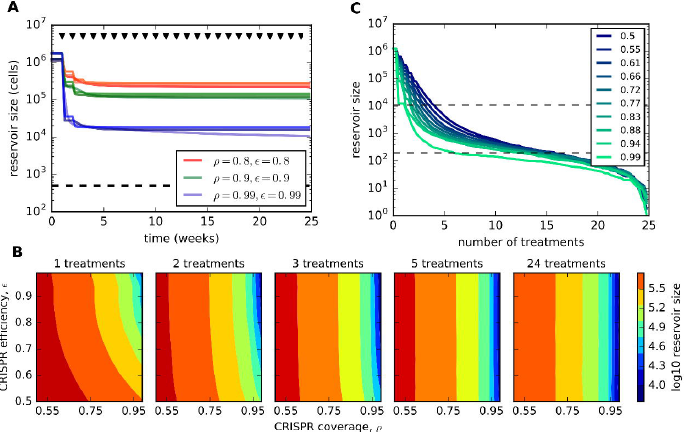
Reservoir depletion with anti-HIV CRISPR therapy. (A) Three representative examples showing the impact of proportional target site conservation (ρ) and enzyme efficacy (ϵ) to a single target site. With a single target site, even at high levels of target site conservation (99%) and a highly efficacious enzyme (ϵ=0.99), reservoir depletion thresholds for functional cure cannot be met. Black triangles indicate dosing and the dashed line represents a stringent threshold for latent reservoir reduction where patients are expected to remain suppressed for years without ART [15]. (B) Simulations with varying CRISPR efficiency and coverage fraction illustrate that after 3-5 hypothetical treatments, reservoir size depletion is constrained predominantly by coverage fraction. Only when coverage is above 95% does reservoir size begin to approach the first clinically relevant threshold of ~10^4^ cells. (C) If 100% coverage of target sites can be achieved (either through multiplexing of targets or due to a target site that is highly conserved), enzyme efficacy becomes relevant, dictating the number of doses to cure. Doses range between 1-5 doses for a median 1 yr remission and 10-15 doses for a potentially lifelong absence of viral rebound based on previously estimated thresholds.

## Discussion

Gene editing using CRISPR/Cas9 has the potential to effect a functional cure for HIV through targeted mutagenesis or proviral genome excision [29]. This approach has now been demonstrated in multiple proof-of-concept *in vitro* and *in vivo* studies [7,9–11,20,30–32]. While laboratory demonstration of gRNA activity has largely relied on clonal populations of lab-adapted HIV strains, clinical applications of this method will need to consider the wide intra- and inter-host diversity of HIV. The global diversity of HIV-1 is reflected in the classification of viruses into four broad groups (M, N, O, and P) that are 25-40% divergent, and within-group subtypes that are up to 15% divergent [23]. This remarkable global diversity of HIV is the result of within-host evolution and adaption to immune pressure, and transmission of genetic variants from the host quasispecies over multiple rounds of viral replication. Target sites chosen for gene editing will therefore also need to reflect this genetic variability within and between individuals.

Globally conserved target sites are good starting points for gRNA design; if their high frequencies in the population are the result of selection, endonuclease-induced mutations are more likely to be highly deleterious to the virus. Indeed, it has been shown that highly conserved target sites are associated with improved antiviral activity, and importantly, delayed viral escape [10,31]. Identification of sites that are conserved at a global or subtype level may also allow for future deployment of these therapies in situations where obtaining individual patient HIV sequence data may not be feasible or practical. To this end, we identified gRNA target sites in HIV LTR that were highly conserved in global consensus sequences and tested the activity of these guides *in vitro*. Using a separate set of deep-sequence data [25], we showed that sites identified from our list of globally conserved targets that were present in the patient’s sequence also showed greater within-host conservation.

Gene therapy approaches designed to cure an infected individual will need to ensure that all relevant within-host variants are targeted. Although early initiation of long-term cART has been shown to reduce the rate of HIV evolution, the virus is still thought to accumulate about 0.97 mutations/kb/year [13,14]. Using a mathematical model, we showed that variants that are not recognized and cleaved will be the major barrier to achieving functional cure thresholds. These variants, if replication-competent, have the potential to reactivate upon cART interruption and reseed the reservoir. Our model makes an assumption about the underlying distribution of quasispecies abundance, which is not fully understood, but notably, three disparate assumptions all resulted in similar conclusions. Estimating time to rebound based on reservoir reduction is challenging and various estimates of thresholds for depletion exist [15,16,33,34]. In our simulations, we have included estimates for median 1 y and median lifetime remission from HIV rebound. However, the depletion itself depends on targeting viral quasispecies diversity, which is not yet fully understood.

The efficiency of gene delivery to target cells is another key factor determining therapeutic outcome. We have also not explicitly incorporated gene delivery in the current model but instead assumed that it is captured within the cleavage efficiency parameter ϵ. However, we have shown previously [35] that gene delivery of endonucleases using viral vectors is prone to large bottlenecks at the points of vector packaging, viral entry and gene expression. Optimization of gene delivery is therefore another important step needed for the clinical success of gene therapies against HIV.

HIV has also been shown to rapidly escape endonuclease targeting [10,11,31], suggesting that therapies will need to target multiple sites concurrently in order to achieve sustained rebound and prevent the emergence of treatment resistance. Our simulations support these findings and show that even enzymes with high on-target efficiency will fail to produce a functional cure if there are target site variants present at frequencies as low as 1%. Two recent proof-of-principle studies showed that an approach with dual gRNAs targeting multiple genes can delay or completely prevent viral escape [30,36]. We identified paired and triplet sets of gRNA target sites that occur in over 98% of the population. Since these sites are likely to also be highly conserved within-host (as our results suggest), they would be good candidates for testing *in vitro* for activity. Although our mathematical model takes multiplexed gRNAs into consideration within the parameter ρ, it does not explicitly include dynamic emergence of treatment-resistant variants. Our model framework is amenable to emergent resistance, but was not included for lack of information on these dynamics. In addition, although many recent studies have targeted LTR, we have shown that *pol* is a better genomic target for directed mutagenesis due to target site conservation. As a result, we believe that targeting multiple sites within *pol* may be a better approach than targeting LTR alone, which generally contains less-conserved sites.

One of the limitations of our within-host analysis is that we do not have detailed information about the patient cohort [25] such as treatment status, age at HIV diagnosis and time of cART initiation and interruption, if any. These factors could potentially impact reservoir diversity. However, the current analysis is primarily aimed at demonstrating the importance and feasibility of designing gRNAs targeting a diverse viral population. Future work needs to address this in greater detail, possibly incorporating treatment-related variables to select gRNA designs.

## Materials and methods

### HIV sequence datasets and pre-processing

For our analysis of global target site conservation, we obtained sequences from the Los Alamos National Laboratory (LANL) database. For each region of interest (*gag*, *pol*, LTR), we downloaded pre-made LANL alignments of all available group M sequences (2016 version). We extracted a majority consensus sequence using Geneious v10 [37] for all sequences in group M and for each subtype.

For within-host analyses of target site conservation, we used deep-sequencing data from a study of HIV-infected blood donors in Brazil [25]. Raw paired-end reads for each patient were trimmed to remove adapters and low-quality regions using Trimmomatic v0.32.2 [38] and mapped using Bowtie2 v0.2 [39] to the consensus sequence deposited by the authors to Genbank. These pre-processing steps (Fig S2) were performed within the Galaxy software framework (https://galaxyproject.org/).

### gRNA target site analysis

We developed a custom script to identify gRNA target sites for an input sequence given a specified PAM sequence (default ‘NGG’ for spCas9) and desired gRNA length *w* (default 20 nt). The algorithm finds all matches to the PAM sequence in the forward and reverse directions and returns, for each match, *w* bases upstream of the PAM sequence. We then used the sgRNA designer from the Broad Institute (https://portals.broadinstitute.org/gpp/public/analysis-tools/sgrna-design) to determine predicted on-target efficacy score and off-target scores (threat matrix) [17]. On-target predicted activity scores are in the range [0,1] with higher values predicting more active guides and a score of 1 indicating successful knockout in the experiments in [17] and [40].

For each target site identified, we determined the number of exact matches found in an alignment of the region of interest (LTR, *gag* or *pol*). We excluded all sites with close off-target matches to the human genome (> 3 matches in Match Bin I, i.e. CFD score = 1 [17]). For each region, we determined pairs and triplets of gRNAs by starting with the previously identified list of gRNAs and adding on guides that increase targeting when used in combination.

We computed target site conservation in terms of the frequency of occurrence of the target site (exact matches) within the alignment and also we used a measure of information content similar to what is used to generate sequence logo plots [41,42]. We applied a moving window of size 23 (corresponding to the width of gRNA) and computed conservation from the relative frequencies of bases in the alignment using the method of Schneider et al. [42] incorporating small-sample correction. The result is a value between 0 and 2 bits with higher values indicating greater sequence conservation. All analyses were performed in R/Bioconductor and code is available on GitHub (http://github.com/proychou/CRISPR).

### Functional testing of gRNA activity

Starting with the list of target sites identified above in LTR, we selected gRNAs from a pool of the top 20 most conserved sites across group M overall, the top 10 most conserved sites in each subtype and the top 20 pairs and triplets. As recommended by sgRNA designer, we excluded any gRNAs with on-target activity scores < 0.2.

We developed 4 LTR-GFP fusion reporter constructs using consensus sequences for all group M, subtype A, subtype B and subtype C (further details in supplement). Internal start codons and stop codons were identified within the sequence for each consensus LTR and the reading frame with the fewest combined number of start codons and stop codons was identified. Reading frame 1 for group M contained 5 start and 4 stop codons, reading frame 1 for subtype A contained 3 start and 6 stop codons, reading frame 1 for subtype B contained 3 start and 6 stop codons, and reading frame 1 for subtype C contained 3 start and 5 stop codons. All the internal start and stop codons were modified for each consensus LTR sequence as follows; ATG to GTG - M to V; TGA to GGA - Stop to G; TAG to GAG - Stop to E; TAA to GAA - Stop to E, so that one continuous open reading frame was generated. Each of the 4 modified consensus LTR sequences was then synthesized as a gBlock and cloned into a reporter plasmid vector (cloning details available upon request) as a fusion to the 5’ end of the eGFP ORF so that the MND promoter drove expression of a single continuous ORF (See Fig S1A for amino acid sequences). The majority of the 59 gRNA target sites identified for analysis within the group M, subtype A, subtype B and subtype C consensus LTRs were not changed by start or stop codon modification, with the exception of overlapping gRNA targets 1 and 2, and overlapping gRNA targets 18 and 19. A separate reporter construct was generated for gRNAs 1, 2, 18 and 19 by fusing their target sequences to the 5’ end of the eGFP ORF so that the MND promoter also drove expression of a single continuous ORF (cloning details available upon request).

Of the 59 LTR-specific gRNA target sites we elected to screen for activity, 23 were present in the group M reporter, 27 were present in the group A reporter, 20 were present in the group B reporter, 18 were present in the group C reporter, and gRNAs 1, 2, 18 and 19 were not present in any LTR-reporter. Three of the gRNA targets were present in all 4 LTR-reporter constructs, 8 were present in 3 LTR-reporter constructs, and 8 were present in 2 LTR-reporter constructs. To screen the activity of individual LTR-specific gRNAs they were cloned into the BbsI site of the plasmid pU6-(Bbs1) CBh-Cas9-T2A-mCherry (a gift from Ralf Kuehn; Addgene plasmid# 64324) under the control of the U6 promoter. This plasmid expresses spCas9 and mCherry from the constitutive CBh promoter. Internal positive controls for GFP knockdown were used by also cloning gRNAs eGFP1 and eGFP2 targeting the sequences CAACTACAAGACCCGCGCCG and GTGAACCGCATCGAGCTGAA into pU6-(Bbs1) CBh-Cas9-T2A-mCherry. To assay gRNA activity 2×10^5^ 293 cells were plated in 12-well plates and the following day individual wells were transfected by PEI transfection with 1000ng of a Cas9/LTR-gRNA expressing plasmid and 250ng of its corresponding LTR-reporter plasmid. At 24 hours post transfection flow cytometry was performed and GFP fluorescence was analyzed in Cas9 expressing (mCherry positive) 293 cells to determine the level of GFP knockdown provided by each gRNA.

### Analysis of flow cytometry data

Raw fcs files were gated using functions from the OpenCyto framework in R/Bioconductor [43] as described previously [35]. Flow data has been uploaded to flowrepository and code is available at http://github.com/proychou/CRISPR.

### Intra-host target site conservation

Focusing on the *pol* gene, we identified spCas9 gRNA target sites within the HIV consensus sequence for each patient using the script described above, excluding any sites containing degenerate bases. We also determined which of the target sites we had previously identified from group- and subtype-level consensus sequences for *pol* were present in the patient consensus sequence. Using the average number of reads overlapping all identified target sites, we excluded any patients with <5000x target site depth since we were interested in variants that may escape targeting by candidate gRNAs. For each target site, we determined the number of reads in the alignment containing an exact match to the target site and excluded any sites where coverage was less than 5000x. We then used the total number of reads that completely overlap the target site to calculate the percentage of exact target site matches.

### Statistical analysis of within-host conservation

To test whether there were differences in target site conservation measured by mean percentages of exact target site matches per total reads, a linear mixed model was fit with percentage as the outcome and the consensus sequence group (group M, subtypes A-C, and patient) as the predictors. A random intercept for each subject by consensus group was used to account for within subject and group variation across the repeated outcomes. An overall test was performed from ANOVA for mixed models using the lmerTest package in R [44]. Post-hoc pairwise tests were also performed comparing the patient-derived sequences, group M, and subtype B (the circulating strain in the patient population). To compare the conservation using patient target sites to the consensus groups, we pooled group M and subtypes A-C into a single group for comparison in the model, while the random effects specification remained the same. P-values corrected for multiple testing were also reported using the Holm method [45].

### Mathematical model of reservoir depletion

We have used a mathematical model of the exponential clearance of the HIV reservoir on consistent ART previously [28]. We extended that model to consider joint treatment with ART and CRISPR gene therapy. Here, the reservoir was modeled as a multi-strain system. For each strain, a clearance rate was chosen from the available data ranges [27] such that the half-life of latently infected cells is normally distributed (indicated by notation N) with mean and standard deviation of 3.6 and 1.5 years respectively, or 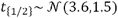. We calculate the clearance rate (per day) for each strain then as 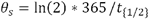. We denote the initial number of latent cells of each strain as where the initial number of cells infected by strain *s* is drawn from a log-normal distribution with average value μ and standard deviation σ = μ/10 so that each strain has size 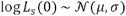. Then, we denote the total number of strains 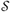 such that 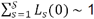 million cells [46,47] and we consider latent reservoirs that begin with average strain sizes μ = 10^3^, 10^4^, and 10^5^ which means that there are 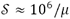 strains, respectively. Then, the model for the size of the reservoir in an individual on suppressive ART undergoing CRISPR treatment is 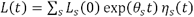

The CRISPR therapy affects some proportion ρ of the strains and has efficiency 㽕. The impact of CRISPR on each strain over time is described by *η_s_*(*t*). For convenience, we model the therapy as a weekly dosage, but can be easily adjusted. Thus, the CRISPR effect is defined mathematically as 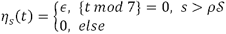. We do not consider the impact of delivery, which was previously described in [35].

The model is simulated using a hybrid stochastic simulation algorithm [48]. When the number of cells in a quasispecies is large (*L_s_*(*t*) > 100), we use the τ leap method [49], and once *L_s_*(*t*) ≤ 100 the simulation proceeds with a direct stochastic “Gillespie” algorithm [50].

## Supporting Information

**Fig S1.** (A) gRNAs were selected for functional testing based on the number of sequences targeted in a global group- or subtype-level alignment either singly, in pairs or triplets (B) Amino acid sequence for the N-terminus of each LTR-reporter GFP fusion construct. M group, subtype A, subtype B, and subtype C reporter amino acid sequences are aligned for each of the 4 reporter constructs. The sequence for eGFP begins with the sequence VSKGEELFT. (C) Transfection efficiency shown in terms of percentage of mCherry+ cells in each treatment. (D) Absolute numbers of mCherry+GFP+ cells in each treatment.

**Fig S2.** (A) Flowchart showing processing steps for intrahost deep sequence data. (B) Target site depth based on number of reads overlapping the target site in an alignment for 4 patients with deep sequence data. Black dots indicate outlier target sites (outside 1.5xIQR) and target sites are grouped and colored according to which consensus sequence they were identified from (the group- or subtype-level consensus from LANL alignments, or from the patient’s HIV consensus sequence).

**Fig S3.** (A) 3 hypothetical distributions of quasispecies abundance in the HIV reservoir. In each case the total size of the reservoir (number of infected cells) is the same(*L* = 10^6^), but the average number of cells in a quasispecies, or “log10 clone size”, is *μ* = 10^3^, 10^4^, 10^5^ respectively. The quasispecies abundances are drawn from a log-normal distribution with variance *μ*/10 in each case. The distribution applies to the simulations in (B) in the same row. (B) Simulations of reservoir decays assuming suppressive ART and hypothetical CRISPR treatment of efficacy *ϵ* and coverage *ρ*. The colored lines indicate quasispecies *L_s_*(*t*), and the solid line shows the total reservoir size *L*(*t*). The dashed line represents a conservative HIV cure threshold taken from the literature. While many quasispecies are removed, the insensitive variants persist and represent a large enough reservoir to prevent cure in all cases.

**Table S1.** (A) Correlation between predicted activity and target site conservation (B) Correlation between measured and predicted activity (C) Correlation between measured activity and target site prevalence

**Table S2.** List of highly conserved, subtype-specific triplet/paired gRNAs

**Table S3.** GFP knockdown with candidate guides tested using fluorescent reporter

**Table S4.** Sequences used in intrahost analysis

**Table S5.** Guides from globally conserved list (using LANL sequences) that have matches in patient sequence

**File S1.** Supplementary methods: LTR reporter design

